# Genomic Analyses of New Genes and Their Phenotypic Effects Reveal Rapid Evolution of Essential Functions in Drosophila Development

**DOI:** 10.1101/2020.10.27.357848

**Authors:** Shengqian Xia, Nicholas W. VanKuren, Chunyan Chen, Li Zhang, Clause Kemkemer, Yi Shao, Hangxing Jia, UnJin Lee, Alexander S. Advani, Andrea Gschwend, Maria Vibranovski, Sidi Chen, Yong E. Zhang, Manyuan Long

## Abstract

It is a conventionally held dogma that the genetic basis underlying development is conserved in a long evolutionary time scale. Ample experiments based on mutational, biochemical, functional, and complementary knockdown/knockout approaches have revealed the unexpectedly important role of recently evolved new genes in the development of *Drosophila*. The recent progress in the analyses of gene effects and improvements in the computational identification of new genes, which has led to large sample sizes of new genes, open the door to investigate the evolution of gene essentiality with a phylogenetically high resolution. These advancements also raised interesting issues related to phenotypic effect analyses of genes, particularly of those that recently originated. Here we reported our analyses of these issues, including the dating of gene ages, the interpretation of RNAi data that may confuse false positive/false negative rates, and the potential confounding impact of compensation and developmental effects that were not considered during previous CRISPR knockout experiments. We further analyzed new data from knockdowns of 702 new genes (~66% of total 1,070 *Drosophila melanogaster* new genes), revealing a similarly high proportion of essential genes from recent evolution, compared to those found in distant ancestors of *D. melanogaster*. Knockout of a few young genes detected analogous essentiality. Furthermore, our experimentally determined distribution and comparison of knockdown efficiency in different RNAi libraries provided valuable data for general functional analyses of genes. Taken together, these data, along with an improved understanding of the phenotypic effect analyses of new genes, provide further evidence to the conclusion that new genes in *Drosophila* quickly evolved essential functions in viability during development.

## INTRODUCTION

The question of how often new genes evolve essential functions is a critical problem in understanding the genetic basis of development and general phenotypic evolution. New genes in evolution have widely attracted discussion (Long and Langley. 1993; Long et al. 2003; Chen et al. 2013; Carvunis et al. 2012; Ding et al. 2013; McLysaght and Hurst. 2015), supported by increasing studies with fulsome evidence in various organisms (e.g. Ruiz-Orera et al. 2018; Xie et al. 2019; Vakirlis et al. 2020; Witt et al. 2019; Jiang and Assis. 2017; Rogers et al. 2014; Schroeder et al. 2020). The detected large number of new genes with unexpected rate of new gene evolution (e.g. Zhang et al. 2019; Shao et al. 2019; Zhang et al. 2010a) and the revealed important functions of new genes (Kasinathan et al. 2020; Lee et al. 2019; Long et al. 2013) challenged a widely held dogma that the genetic basis in control of development is conserved in a long time scale of evolution (Ashburner et al. 1999; Gould. 2002; Carroll. 2005; Krebs et al. 2013). Our previous work used the RNAi knockdown in a smaller sample showing that new genes may quickly become essential in *Drosophila* and that potential for a gene to develop an essential function is independent of its age (Chen et al. 2010). This work suggests a tremendous and quickly evolving genetic diversity, which had not been previously anticipated. Since then, genomes of better quality from more species have allowed for more reliable new gene annotation (Shao et al. 2019). In addition, technical progress in the detection of gene effects has increased with better equipped knockdown libraries and direct CRISPR knockout methods. Related scientific discoveries and technical development in knockdown and knockout techniques -- e.g., Green et al (2014) and Kondo et al (2017), respectively -- can be considered when investigating the evolution of gene essentiality.

We will present in this report our recent experiments and computational analyses, examining a few important issues raised in recent years (e.g. by Kondo et al. (2017) and Green et al. (2014)) that we find to be generally relevant for the detection of the phenotypic effects of genes, particularly of those that recently originated. Our investigations include the following: 1) the estimation of new gene ages; 2) an evaluation of the knockdown efficiency distribution in RNAi experiments; 3) an understanding of the differences between different RNAi libraries in phenotyping large samples of new genes for viability effects; 4) an interpretation of knockout data regarding the compensation effect. Our analyses, with additional evidence published recently by our group and others, provide ample and strong evidence to further support a notion suggested by the fitness effect analysis of new genes in *Drosophila*: new genes have quickly evolved essential functions in viability during development.

## RESULTS AND DISCUSSION

### Identification of *Drosophila*-specific genes by the age dating

Two new gene datasets are available for *D. melanogaster*, which include the dataset of Kondo *et al* (2017; the K-dataset, the underlying pipeline as the K-pipeline) and the dataset we recently reported (Zhang et al, 2010b; Shao et al, 2019 called as the GenTree Fly dataset, the G-dataset). In order to determine which dataset is more accurate and thus could be used in the downstream analyses, we estimated their qualities by performing systematic comparison. Kondo *et al*. (2017) identified 1,182 new genes that postdated the split of *D. melanogaster* and *D. pseudoobscura* (Branches 3~6, ~40 Mya(million years ago); Fig. 1A and 1B). They inferred the ages of these genes by incorporating the UCSC DNA-level synteny information, homology information based on comparison of annotated proteins and RNA expression profiling. By contrast, we identified 654 new genes in this same evolutionary period (Fig. 1A and 1B) using the same UCSC synteny information (Rhead et al. 2010) and our maximum parsimony-based pipelines (Zhang et al. 2010b).

**Figure 1.**
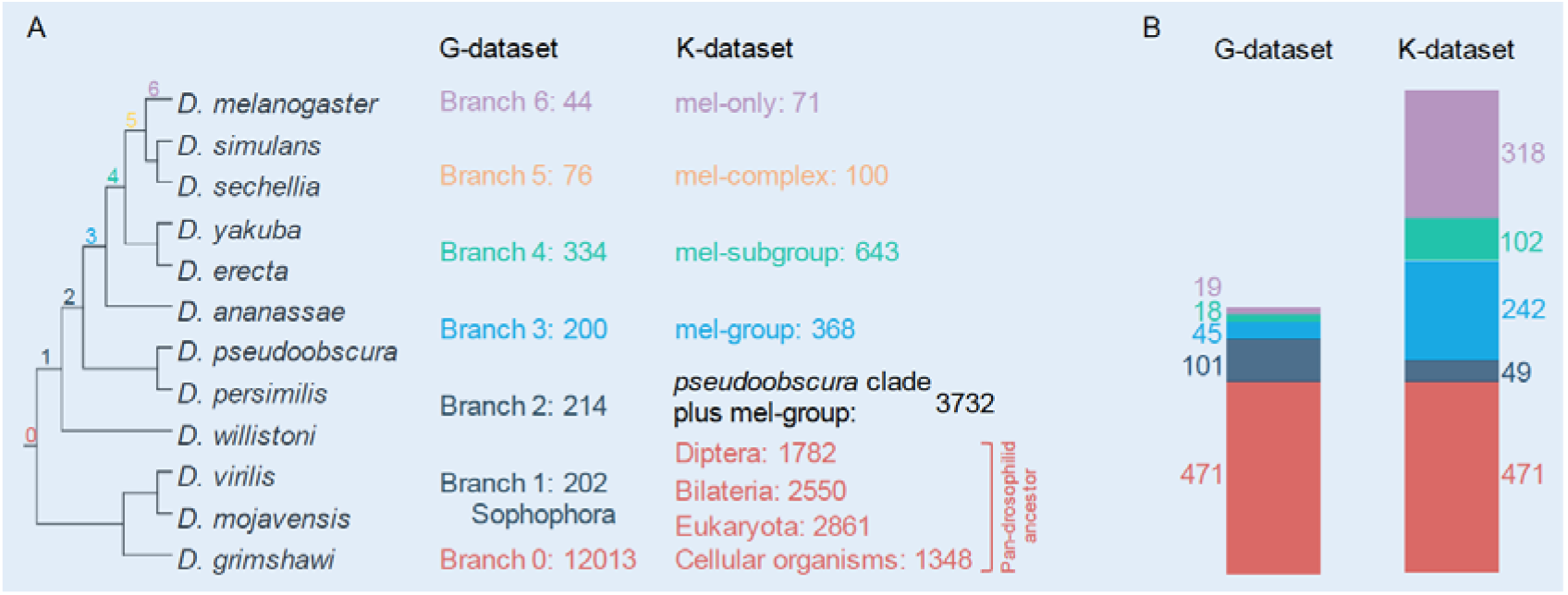
Summary of new gene candidates in the K-dataset and G-dataset. A. phylogenetic distribution of gene origination identified by the K-pipeline and the G-pipeline as shown in the two datasets. B. Evaluation of the two datasets based on individual gene analyses. The two datasets share 471 candidates (red). The G-dataset consist of 101 authentic candidates (deep blue) undetected in the K-dataset, 19 false positives (light purple), 18 dubious cases (green) and 45 cases not applicable for dating (sky blue). By contrast, the K-dataset includes 49 bona fide new gene candidate, 318 false positives, 102 dubious cases and 242 difficult cases. Note, the K-dataset mentions 1,182 genes in the main text, however its associated supplemental table includes 1,176 genes with 6 genes listed more than once.

We investigated why the K-dataset was almost twice the G-dataset over the same evolutionary period. The K-dataset contained 471 new gene candidates in the G-dataset (471/654 = 72%) (Fig. 1B, red), among which 313 are of the exact same ages while 158 show either younger (123 genes) or older (35 genes) (Supplemental Table S1). For the remaining 183 genes absent in the K-dataset, we found 101 as authentic new genes (Fig. 1B, deep blue; Supplemental Table S2) after extensive validation by manually checking UCSC synteny information and four additional resources including the FlyBase ortholog annotation, Ensembl Metazoa homolog annotation, protein prediction in outgroup species and published literatures (see also Materials and Methods). This result indicates a high false negative rate in the K-dataset. By contrast, only 19 genes are old genes, which represent false positives in the G-dataset (Fig. 1B, light purple). Then, for 45 genes (Fig. 1B, sky blue), they are located in clusters of tandemly amplified genes or transposon-rich regions, where synteny is often ambiguous or difficult to build especially when outgroup species genomes are poorly assembled. Their ages are difficult to infer. Analogously, the final remaining 18 candidates (Fig. 1B, green) are dubious, either with inconsistent topology between UCSC synteny information and Ensembl tree, marginal protein similarity between species or gene structure model changes.

We next examined the 711 (1182-471=711) new gene candidates unique to the K-dataset by manually examining phylogenetic distribution and their syntenic relationship with genes in various species. We could only confirm 49 authentic new genes, which represent false negatives of the G-dataset. By contrast, 318 out of 711 genes were incorrectly dated as new genes due to four problematic practices (Fig. 1B; Supplemental Table S2 lists the 318 false positives): 1) neglecting 275 that have orthologs in outgroup species; 2) taking 32 noncoding or pseudogene models as protein coding genes; 3) treating 6 redundant entries of same genes as different genes; and 4) misdating 5 polycistronic coding genes reported by the literatures. In addition, 242 (Fig. 1B, sky blue) genes are located in repetitive regions. To be conservative, the G-dataset excluded these genes from dating. Then, the remaining 102 candidates (Fig. 1B, green) are dubious.

We conclude, based on above exhaustive manual evaluation, that the G-dataset is of much higher quality compared to the K-dataset: 1) the false negative rate and false positive rate of the G-dataset is estimated as 7.5% (49) and 2.9% (19), respectively; 2) The both parameters of the K-dataset are higher, 8.5% (101) and 26.9% (318), respectively; 3) the G-dataset only contains 9.6% (63) low-quality candidates (genes in repetitive or dubious categories), while the K-dataset consists of 29.1% (344) such candidates. Overall, 56% (662/1,182) new gene candidates in the K-dataset are either false positive or dubious.

### Measuring reproducibility and efficiency of knockdown

We investigated the consistency of RNAi experiments with the same lines and the same drivers in different laboratories, conditions, and years. Zeng et al. (2015) screened 16,562 transgenic RNAi lines using an Act5C-Gal4 driver to detect the lethality of 12,705 protein-coding genes (~90% of all annotated coding genes) in their study of intestinal stem cell development and maintenance. Their dataset included RNAi lines targeting the same 103 genes that were measured for lethality by Chen et al. (2010). Chen et al. (2010) and Zeng et al. (2015) obtained the same phenotypes for 88 (85.4%) genes, including 30 (29.1%) of the lethal phenotype and 58 (56.3%) of non-lethal phenotype (Fig. 2A, Supplemental Table S3). These data suggest that despite differences in independent observers, lab environments, and years to conduct experiments, the vast majority of RNAi knockdown experiments are reproducible for phenotyping lethality and non-lethality.

**Figure 2.**
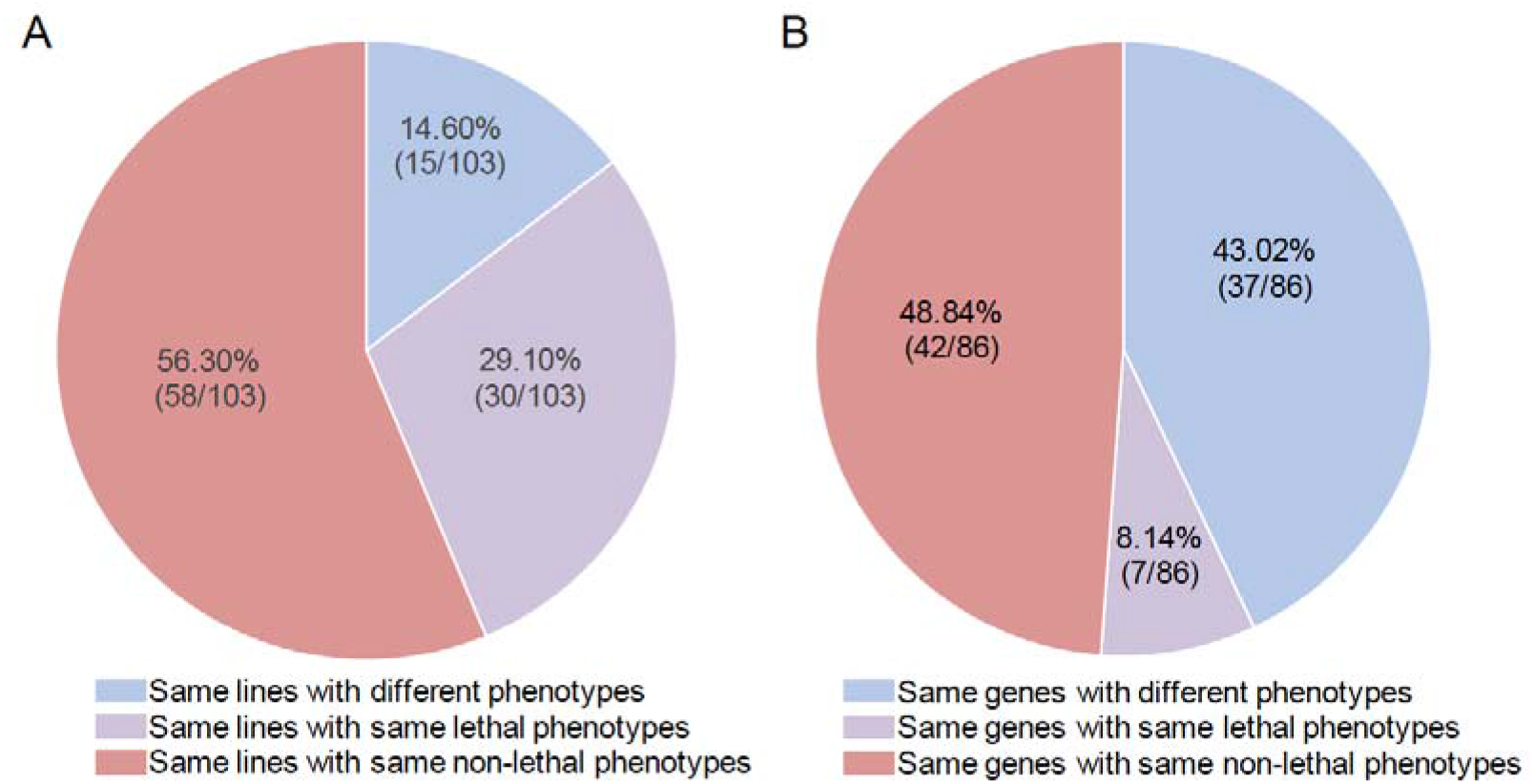
The reproducibility analysis of RNAi experiments by comparing two groups of independent experiments by Chen et al (2010) and Zeng et al (2015). A. Phenotypes of same 103 RNAi lines analyzed by Chen et al (2010) and Zeng et al (2015) using same lines; B. Phenotypes of 86 same new genes knocked down by two different drivers or the same drivers with different insertion sites. The old drivers detected 29 genes as lethal while 57 non-lethal; the new drivers detected 20 genes as lethal while 66 non-lethal.

We also tested consistency between RNAi lines with different RNAi drivers (called new drivers) or same drivers in different genome positions. Specifically, the datasets of Chen et al (2010) and Zeng et al (2015) shared 86 new genes in knockdown experiments, mostly (81.4%, 70) with different RNAi drivers and fewer (18.6%, 16) same drivers in different genome positions (Supplemental Table S4). This dataset showed that: 7 genes were consistently lethal; 42 genes were consistently non-lethal; and 37 genes have different phenotypes (Fig. 2B). Thus, the two groups with different drivers or same drivers with different positions show that more genes (57.0%, 49) have the same phenotypes.

We considered an additional factor in RNAi knockdown, sensitivity, in the two widely used RNAi libraries: the Vienna Drosophila Resource Center’s (VDRC’s) GD and KK libraries (Dietzl et al., 2007). The GD libraries were constructed using P-elements to randomly insert hairpin RNAs (average 321bp) into the genome targeting individual genes, while the KK library inserted constructs carrying hairpin RNAs (average 357bp) into a specific landing site by ΦC31-mediated homologous recombination. All KK lines carry an insertion at 30B3, but a proportion (23-25%) also carry an insertion at 40D3 (*tio* locus) that results in pupal lethality when using constitutive drivers like Act5C-GAL4 (Green et al. 2014; Vissers et al. 2016). Unless specified, no lines discussed below contain 40D3 insertions.

Given the intrinsic different designs of GD and KK libraries, we hypothesized that they have different false negative or false positive rates, which cause the inconsistency shown in Fig. 2B. Only GD lines were examined previously, and they have a high false negative rate (39.9%) but low false positive rate (<2%) (Dietzl et al. 2007). The high false negative rate is likely caused by insufficient target gene knockdown, while false positives may be due to off target effects (Dietzl et al. 2007). We thus tested the knockdown efficiency of 75 KK lines targeting randomly selected 75 young genes (Supplemental Table S5, Fig. 3A). We found that the knockdown efficiency of KK lines is generally lower than the efficiency of 64 GD lines as previously reported (Dietzl et al. 2007). Specifically, using the same driver (Act5C), we found that in general, GD lines have significantly higher knockdown efficiency than KK lines, as shown by the knockdown expression as the percentage of the control expression (Fig. 3A). That is, the KK lines have an average knockdown efficiency as 48.6% of control expression while the GD lines show an average efficiency as 38.1% (Fig. 3B and 3C, *t*-test *P* = 0.031). Notably, the expression reduction to 50~60% level of the wide-type level was observed to have no significant fitness loss due to widespread haplosufficiency (Huang et al. 2010; VanKuren and Long, 2018). Detecting any fitness effect may be expected when the expression drops to a lower level, for example, 20~30% or lower of the control expression. In this range of knockdown efficiency, we observed that only 29% of KK lines but 41% of GD lines reduced target expression levels to ≤20% of control levels; 37% of KK lines but 53% of GD lines were seen to reduce target expression levels to ≤ 30% of control levels (Fig. 3A). Thus, it is expected that GD lines have a significantly higher power in detecting lethal phenotypes as shown in the next section.

**Figure 3.**
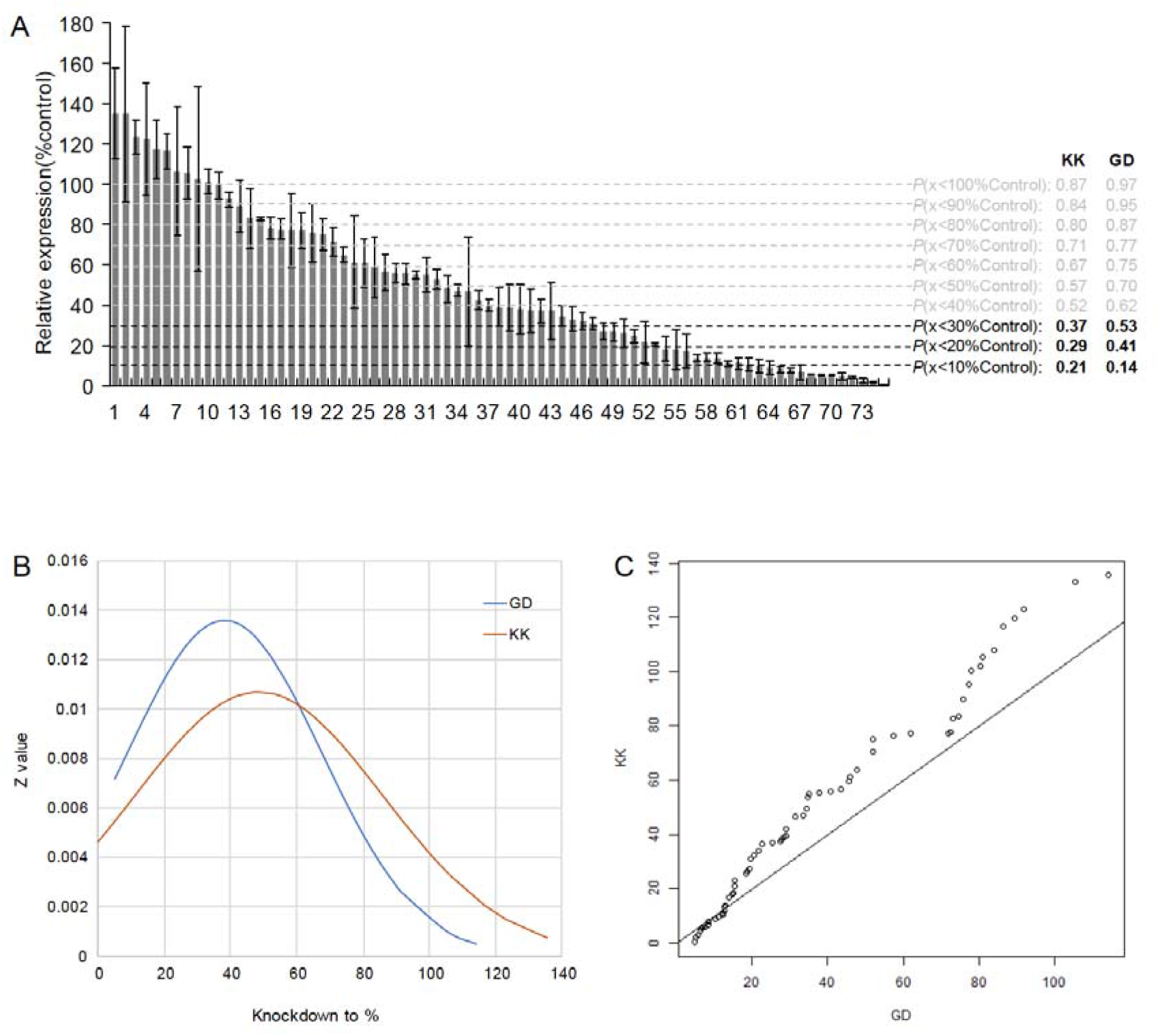
Knockdown efficiency in the KK and GD libraries revealed GD lines have significantly higher knockdown efficiency than the KK lines. A. The knockdown efficiency of the 75 KK lines was measured, compared to the expression of the wild-type control and the standard deviation is calculated from the measurement of three repeats; *P* refers to proportion of genes with the expression lower than a certain threshold while the values of KK lines are generated in this work and that of GD lines are extracted from Dietzl et al. (2007). B. The distributions of knockdown efficiency of KK and GD lines. C. The Q-Q Plots between KK and GD lines.

To estimate false positive rate of KK lines, we constructed 10 randomly chosen new KK lines targeting one member of a young duplicate gene pair, in addition to one KK lines and 3 TRiP lines (Transgenic RNAi Project, BDSC, Materials and Methods). The rationale is that for each gene of interest its paralog is the most likely off target. The same rationale was also followed by (Dietzl et al. 2007) when false positive rates of GD lines were estimated. We measured the knockdown efficiency and estimated off-target effects using these 14 lines with qPCR experiments in adult whole bodies (Fig. 4). We found that two lines likely produce off-target effects (*NV-CG31958, 34008* (the TRiP line)), for both of which the expression of paralog is down-regulated to similar or even lower level compared to the corresponding gene of interest. Twelve other lines have significantly higher target effects than off-target effects, among which 10 genes reduced activity to 20-80% expression level of the control (7 genes reduced activity to 20-40%) and only two genes (*CG32164, CG7046*) reach≤20% of control levels. Thus, if we take 20% as the cutoff of efficient knockdown, only *CG31958* could be counted as the false positive, and *CG32164* and *CG7046* be counted as the true positives. Collectively speaking, the off-target effects are rare while insufficient knockdowns are pervasive.

**Figure 4.**
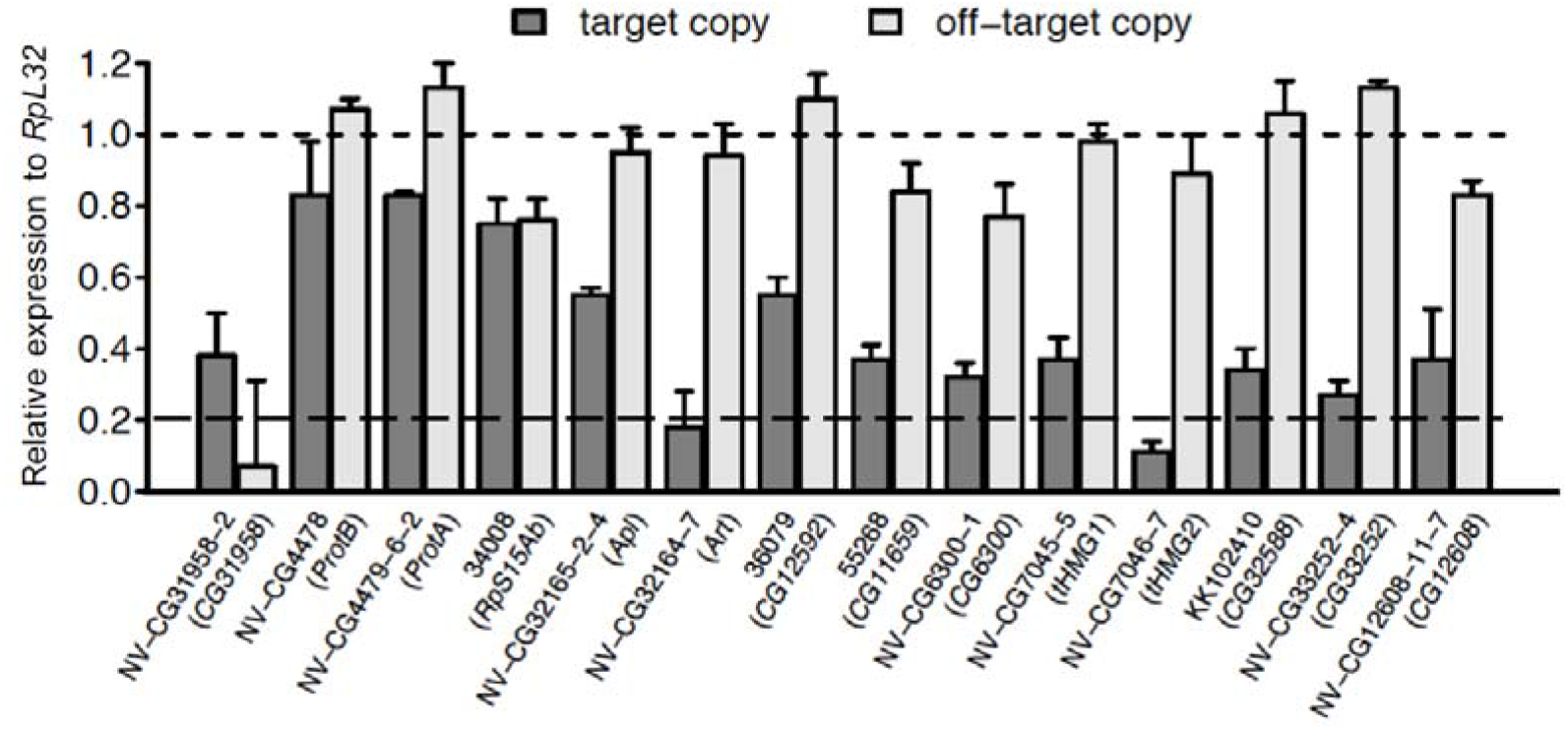
Experimental comparison of the efficiency and off-target effects explain the conservative nature of RNAi knockdown experiments and limited off-targets propensity. For each young duplicate gene pair specific for *D. melanogaster* and *melanogaster* species complex, we examined their expression intensity relative to the wide type control in whole body flies with qPCR. The standard deviation is calculated based on three replicates.

These experiments detected a variation of knockdown efficiency among different drivers where newer KK lines have lower efficiency and thus higher false negatives compared to older GD lines. Therefore, these observations offer an alternative interpretation of the incongruence than the false-positive-only rationale of Kondo et al (2017): when new RNAi drivers were added to the analysis, insufficient knockdown was also introduced with a high probability. This would create incongruence between old and new drivers if the old and new triggers have significantly different sensitivity.

### Phenotyping essentiality of new genes in RNAi libraries

We first investigated differences in measured lethality between the KK and GD libraries used in Chen et al (2010). To control for the confounding effect of *tio* insertion in the KK lines, we genotyped these lines using PCR-amplification and found that out of 153 KK lines we collected, 47 (30.7%) had two landing sites and 6 (3.9%) had only 40D3 landing site (the confounding site) (Green et al. 2014), which all showed lethal phenotypes (Supplemental Table S6). Using the recombination approach (Green et al. 2014), we recovered 41 of the 47 lines into the lines that have only the 30B3 site. The RNAi knockdown of 140 KK lines carrying insertions only at 30B3 identified 12 genes (8.6%) with lethal phenotypes (Supplemental Table S6). Meanwhile, 12 genes in 59 GD lines (20.3%) were detected to have lethal knockdown effects (Chen et al. 2010), significantly higher than the KK libraries (*P* = 0.0112, Fisher’s Exact Test). As aforementioned, this difference is likely due to higher false negative rate of KK lines (Fig. 3).

By using the essentiality data of 10,652 old genes provided by VDRC (https://stockcenter.vdrc.at/control/library_rnai) that were in branch 0 (Shao et al. 2019), we characterized the statistical distribution of essential old genes (Fig. 5). We independently sampled 1000 times, with each randomly sampling 150 old genes and calculating the proportion of essential ones. We found that in the GD library, the probability to obtain a proportion of essential new genes equal or lower than 20.3% is 0.780. Meanwhile, in the KK library, the probability to observe a proportion of essential new genes equal or lower than 8.6% is 0.867. These analyses of GD and KK libraries reveal similarly that the proportions of new and old genes with lethal phenotypes are not statistically different.

**Figure 5.**
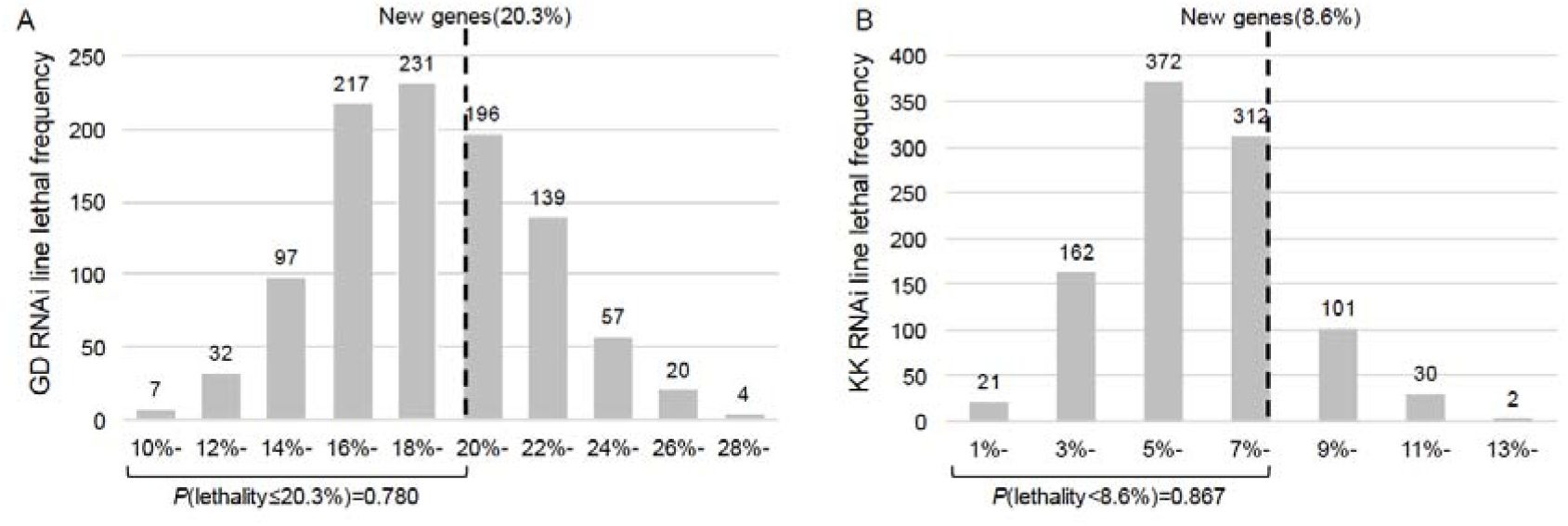
Comparison of proportions of lethality between new genes and old genes in GD lines (**A**) and KK lines (**B**) suggests that in both GD and KK lines, new genes have an equally high probability to be lethal as old genes. Since old genes are much more abundant than new genes, we generated 1000 random sample of old genes with the same number of new genes and then plotted the distribution of proportion of essential genes as histograms.

Further analysis of gene essentiality data in a recent version of VDRC libraries (retrieved online in April 2019) detected with increased resolution the proportions of essential genes in six detectable ancestral stages of *D. melanogaster*. We reported the analysis of the GD library, which has a significantly higher knockdown efficiency than the KK library. In total, 11,354 genes (72% of 15,682 genes in the species, Ensembl 73) have been phenotyped for their lethality or nonlethality, including 702 *Drosophila* genus specific genes (66% of 1,070 detected *Drosophila*-specific genes) (Long et al. 2013; Shao et al. 2019) and 10,652 genes that predated the *Drosophila* divergence 40 Mya.

We parsimoniously mapped the 702 *Drosophila*-specific genes on the six ancestral stages by examining their species distribution in the *Drosophila* phylogeny (Shao et al. 2019) (Fig. 6A). Of the 702 genes, 19.7% (138) are directly observed to be essential, similar to the proportion of essential old genes, 18.9% (*P* = 0.6212, Fisher’s exact test). We considered a low knockdown efficiency as shown by the 47% of GD lines whose knockdowns are expressed at the level of 30% or higher of the control (Fig. 3A), suggesting that 47% of RNA lines are invalid for the testing and should be subtracted from the total tested lines.

**Figure 6.**
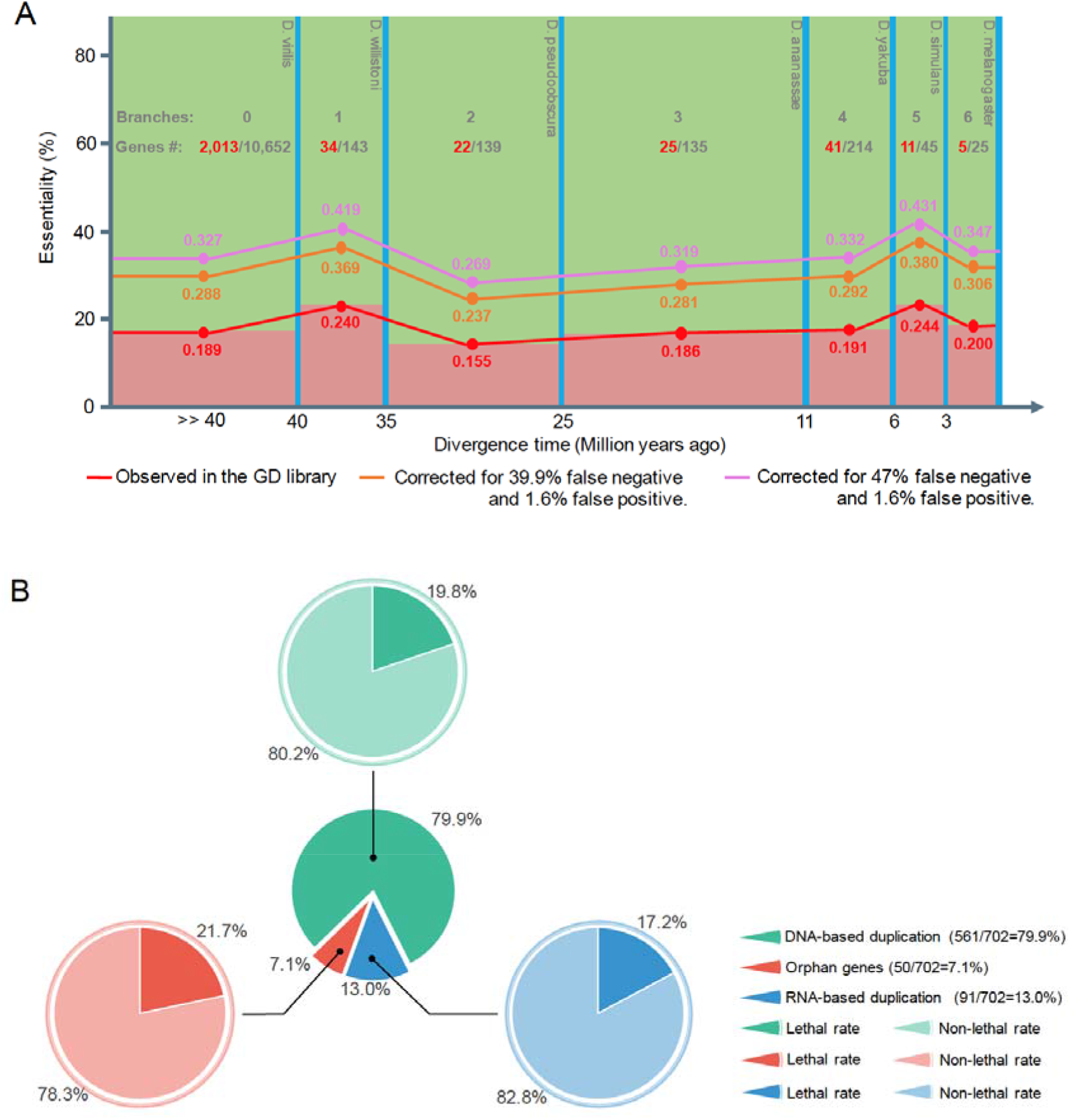
Lethality proportion of 702 *Drosophila*-specific genes. A. Lethality proportion of 702 *Drosophila*-specific genes in 6 ancestral stages of extant *D. melanogaster*, compared to the lethality proportion of 10,652 genes older than 40 Mya. No stages show an essentiality proportion significantly different from that of old genes (0.189). B. Lethality proportion of 702 *Drosophila*-specific genes based on three origin mechanism catalogs. No catalog shows a lethality proportion significantly different from that of old genes (0.189).

Thus, the actual proportion of essential genes can be estimated by correcting for the bias of false positives (Fp) and false negatives (Fn) by following formula:

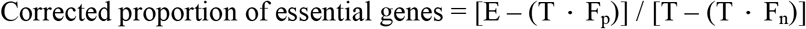

Where E and T are observed number of essential genes and total number of genes examined, respectively. F_p_ was measured as 1.6% (Dietzl et al., 2007) while F_n_ as 47% as estimated above or 39.9% as measured previously (Dietzl et al., 2007). Thus, the estimated proportion of essential genes after correcting false positives and false negatives can be as high as 36.5% for the estimated false negative rate of 47% in this study. The corrected proportion can be also as high as 32.2% given the previously measured false negative rate of 39.9%. Furthermore, all six stages show a stable proportion of essential genes; none of the proportions is statistically different from the proportion of old genes (Fig. 6A). Meanwhile, lethal rates of new genes which belong to three origin mechanism categories (DNA-based duplication, RNA-based duplication and orphan genes, Shao et al., 2019) also show no significant difference (Fig. 6B). Interestingly, 21.7% of orphan genes, some of which might be *de novo* genes (Long et al., 2013), are essential. These data add new insight into the evolution of essentiality in all ancestral stages: soon after genes originated and fixed in *D. melanogaster*, a stable proportion of new genes is essential throughout entire evolutionary process from ancient ancestors to the speciation of *D. melanogaster*.

These data of knockdown experiments on a large number of new genes further supported what we proposed before: *Drosophila* new genes rapidly evolve essential functions within the divergence of *Drosophila* genus; knockdown of these genes leads to death of flies.

### Analyses of mutants identified young essential genes

Kondo et al (2017) recommended and used CRISPR/Cas9-mediated mutagenesis to create small frameshift indel mutations in targeted genes. This method has two potential issues. First, it is now well documented that vertebrate cells such as mammalian cells or zebrafish cells recognize such aberrant mRNAs and compensate for their loss by increasing expression of genes with high sequence similarity, such as paralogs in zebrafish, worm and other organisms (Rossi et al. 2015; El-Brolosy and Stainier 2017; El-Brolosy et al. 2019); Ma et al, 2019; Serobyan et al, 2020). This has the effect of producing false negatives especially for recent duplicates. We confirmed that a similar compensation effect exists in *Drosophila*. Specifically, when we induced a one-nucleotide deletion using CRISPR/Cas9 into the ORF region of *vismay* (*vis*), a *D. melanogaster*-specific gene duplicated from a parental gene, *achintya (achi*), 0.8 Mya, with a nucleotide similarity of 92% between the two copies. We found that *achi* in the *vis* mutant was significantly upregulated whereas a randomly selected unrelated gene *CG12608* and the distantly related gene *hth* (nucleotide similarity of 45%) to *vis*, did not show such an effect (Fig. 7). Second, CRISPR/Cas9-mediated mutagenesis cannot detect the effects of maternal and paternal effect genes, which can be common in *Drosophila* (Perrimon et al. 1989; Raices et al. 2019) and can be detected by RNAi knockdown. Therefore, the two approaches of knockdown and knockout/mutagenesis are complementary to each other given their technical characteristics.

**Figure 7.**
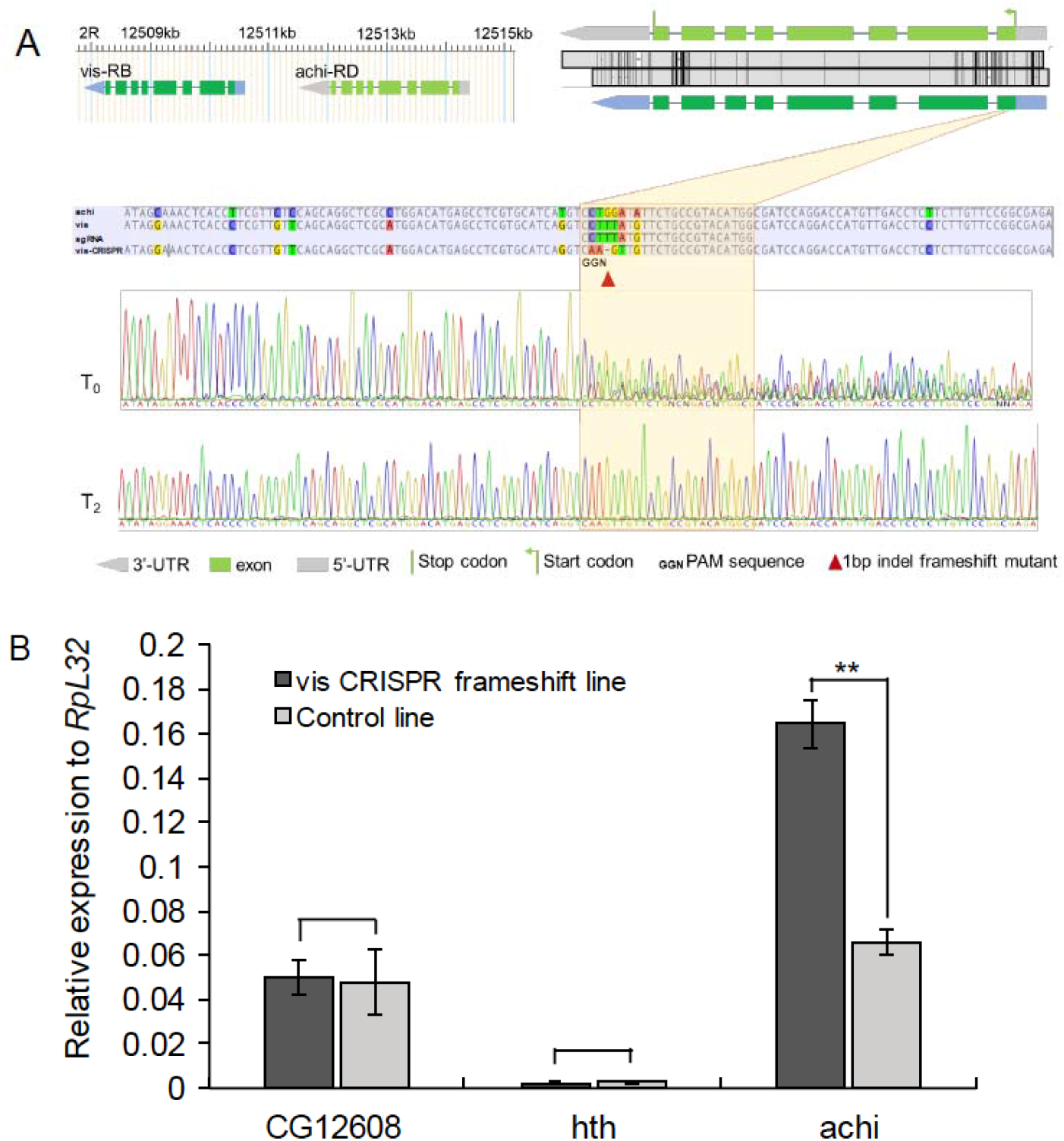
CRISPR/Cas9 frameshift mutant could induce compensatory effect in *Drosophila*. A. Design of CRISPR/Cas9 mutant. We targeted a randomly chosen young gene, *vis*, which emerged via duplication of *achi* in the common ancestor of *melanogaster* species complex. The genomic arrangement of two genes are shown in the upper left panel with the boxes referring to exons and connecting lines as introns. The pair shares a high sequence identity (0.92) in their 9 exons, which is schematically shown in the upper right panel. The middle panel shows the diverged site between *vis* and *achi*, which was chosen to design a short guide RNA (sgRNA) specifically targeting *vis*. The mutation (CTTTA→AAGT) was marked with a red triangle. The raw sanger sequencing data for the initial generation (T0) and the second generation of offspring (T2) was shown. B. The compensation effect of *achi*. In the frameshift mutant of *vis, achVs* expression issignificantly increased (*P* = 0.0003). By contrast, the unrelated *CG12608* and the remotely related *hth* did not show any significant upregulation. *RpL32* was used as a control as in (VanKuren and Long 2018).

Actually, in-depth analyses of several cases already provided further evidence supporting essentiality of new genes in development. First, Ross et al (2013) reported a stepwise neofunctionalization evolution in which a centromere-targeting gene in *Drosophila, Umbrea*, was generated less than 15 Mya. Both RNAi knockdown, rescue experiments and P-element mediated gene knockout revealed that *Umbrea* evolved a species-specific essentiality to target centromere in chromosome segregation (Chen et al. 2010; Ross et al. 2013). Second, Lee et al (2019) recently detected stage-specific (embryos/larvae/pupa) lethality associated with RNAi knockdown and CRISPR knockout in *Cocoon*, a gene emerged 4 Mya in the common ancestor of the clade of *D. melanogaster*-simulans. These data show that *Cocoon* is essential for the survival at multiple developmental stages, including the critical embryonic stage. Third, P-element insertion/excision experiments show the essentiality of *K81* as a paternal element in early development. This gene only exists in the *Drosophila melanogaster-subgroup* species that diverged 6 Mya (Loppin et al. 2005). Fourth, Zeus, a gene that duplicated from the highly conserved transcription factor *CAF40* 4 Mya in the common ancestor of *D. melanogaster* and *D. simulans* rapidly evolved new essential functions in male reproductive functions, as detected in the null mutants and knockdown (Chen et al. 2012; Ventura, 2019). Fifth, A pair of extremely young duplicates, *Apollo* (*Apl*) and *Artemis (Arts)*, was found to have been fixed 200,000 years ago in *D. melanogaster* populations (VanKuren and Long, 2018).

CRISPR-created gene deletions of these genes showed that both evolved distinct essential functions in gametogenesis and *Apl* critical function in development. Sixth, in a comprehensive functional and evolutionary analysis of the ZAD-ZNF gene family in *Drosophila* (Kasinathan et al, 2020), 86 paralogous copies were identified with phenotypic effects detected by knockdown and knockout in *D. melanogaster*. It was found that the proportion (17/58 = 29.3%) of lethal copies in old duplicates (>40 Mya) and the proportion (11/28 = 39.3%) of lethal copies in *Drosophila*-specific duplicates (<40Mya) are statistically similar. Further functional analyses of one of the non-essential young copies (*CG17802, Nicknack*) reported by Kondo et al (2017) clearly unveiled an essential function for larval development. These pieces of evidence strongly support the notion that new genes can quickly evolve essential functions in development.

### Concluding Remarks

We appreciate the extensive experiment, computation and data-compilation by Kondo et al (2017) and their interests in the evolution of gene essentiality. However, we found that the K-pipeline and K-dataset were associated with a high false positive rate. Moreover, their interpretation of RNAi data is problematic due to confusing the false negative and false positive, while they applied an incorrect CRISPR mutagenesis that neglected compensation and parental effects. The data we created in this study, while revealing their errors and technical insufficiencies, increase understanding of technical subtleties for analyzing effects of young duplicate genes and other genes. More data and additional analyses of related scientific issues for the testing of fitness and functional effects of new genes from the two complementary approaches, RNAi knockdown and CRISPR knockout, provided a strong support for the concept: the new genes rapidly evolved essential functions in development in *Drosophila*. This challenges a conventional belief in the antiquity of important gene functions in general (Jacob, 1977; Mayr, 1982; Ashburner et al. 1999; Krebs et al. 2013) and in development process in specific (Gould. 2002; Carroll. 2005).

## MATERIALS AND METHODS

### Comparison of the K-dataset and the G-dataset for Drosophila new genes

#### Overall comparison scheme

To our knowledge, there is no published genome-wide evaluation of gene ages in *Drosophila*. Specifically, around a decade ago, we took advantage of the syntenic genomic (DNA) alignment generated by the UCSC group and performed the genomewide age dating for the first time in *Drosophila* (Zhang et al. 2010b). At that moment, we compared our data to previous studies based on limited number of cases and discovered the general reproducibility across studies. The genome-wide dataset by Kondo *et al*. enabled a systematic large-scale comparison. We began with repeating our pipeline on FlyBase annotation v6.02 and identified 654 new genes originated on the branch toward *D. melanogaster* after the species split of *D. melanogaster* and *D. pseudoobscura* (Fig. 1). Concurrently, based on similar FlyBase release v6.13, Kondo *et al*. performed dating according to the same syntenic alignment of UCSC, which is further complemented by v6.02 protein-level BLAST search and filter with testisspecific expression (Kondo et al. 2017). With additional Dollo-parsimonious searches, they identified 1,182 new genes originated in the same period including *melanogaster*-group to *melanogaster-only*, which correspond to age group 3 to 6 in our analysis, respectively (Supplemental Fig. S1). Since the FlyBase annotation version is similar (v6.02 vs. v6.13), only 12 entries out of the G-dataset and K-dataset are not comparable due to “Gene model change” (Supplemental Fig. S1). They represent either expired models or new models in v6.13. Except them, all other genes can be compared across two datasets.

We found that 471 out of the 654 new gene candidates in the G-dataset are covered in the K-database by comparing the Ensembl IDs of these databases. Moreover, 313 (66%) genes show the exact same ages. Since manual curation needs extensive efforts, we did not examine why the remaining 158 genes show minor age difference. Instead, we subsequently only focused on those genes which show conflicting dating results, *i.e*., included or excluded in new gene dataset across two studies. As a result, we classified the conflicting cases into six major categories, which can be further divided into around 20 more specific sub-categories (Supplemental Fig. S1). We documented how we performed classification as below.

#### Four independent information sources facilitate evaluations of two age datasets

The challenges in the dating of gene age largely lie in the ambiguity of calling orthologs across outgroup species (Liebeskind et al. 2016). We found that the conflict of age dating was often due to the difference of DNA-level synteny and protein-level homology search. Specifically, for a gene of interest, *A*, the UCSC best-to-best synteny information shows that its ortholog is present in one outgroup species, *B*. However, the protein-level information may reveal an absence. The opposite scenario can occur too. In these conflicts, we turned to independent resources including FlyBase ortholog annotation, the homolog annotation and gene family tree provided by Ensembl Metazoa (St Pierre et al. 2014; Kersey et al. 2015), protein prediction in outgroup species based on gene models of *D. melanogaster* and literatures. Specifically, FlyBase provided AAA (Assembly/Alignment/Annotation) syntenic ortholog annotation. If species *B* encodes a FlyBase annotated ortholog, gene *A* likely predated the species-split of *D. melanogaster* and *B*. Similarly, Ensembl Metazoa provided one-to-one best-to-best ortholog annotation. We used it like FlyBase. Finally, for some cases where synteny predicted orthologous regions of *B* do not harbor an annotated gene, we conceptually translated this region with the protein of *D. melanogaster* as the template. BLAST (Tblastn) was used here. We have two reasons to perform additional annotation: 1) recently evolved genes are often poorly annotated; 2) annotation quality of outgroup species is presumably worse compared to *D. melanogaster* and we need to correct this bias. For particularly interesting cases (*e.g*. polycistronic coding genes), we searched literatures describing their evolutionary history.

#### Conflicting cases could be classified into six categories

We implemented a series of customized rules to call the presence of ortholog of gene *A* in species *B*. The first set of rules are used to call presence of ortholog based on gene prediction. For a synteny-predicted candidate orthologous region in *B*, we ran Tblastn to predict whether this region encodes an orthologous protein of *A*. If Tblastn could align the protein of *D. melanogaster* beyond the following thresholds (identity cutoff > 70% & coverage cutoff > 30%, identity cutoff > 30% & coverage cutoff > 70%, identity cutoff > 50% & coverage cutoff > 50%), we believed that the ortholog is present. If the alignment meets with the threshold (identity cutoff < 30% & coverage cutoff < 30%), the ortholog of *A* is absent in *B*. For all other cases, we called them as “boundary cases” if there is also no ortholog annotated by FlyBase and Ensembl. A total of 80 candidate new genes fall in this category including 65 cases in the list of Kondo *et al*. and 15 cases in our list (Supplemental Fig. S1).

Secondly, for 275 out of 318 new genes dated by Kondo *et al* but not by us (Supplemental Fig. S1), we identified orthologs as supported by at least two independent sources (FlyBase, Ensembl homolog, Ensembl gene family tree, and/or prediction). For example, in case of *FBgn0027589*, the ortholog is present across all 12 *Drosophila* species, which is supported by both Ensembl and FlyBase. The remaining 43 new genes are misidentified due to other types of problems (*e.g*. annotation problem due to polycistronic structure such as *tal-1A/tal-2A/tal-3A*) All these 318 genes are marked as “Dating problem in Kondo *et a?* (Supplemental Fig. S1). In the opposite scenario, 101 new genes are only identified by us, which could be divided into four cases: 1) for 50 genes called as old genes in the K-dataset, their new gene calling were also supported by lack of one-to-one orthologs annotated by Ensembl or FlyBase; 2) for 5 genes called as old genes in the K-dataset, they are subject to complex evolutionary trajectories (pseudogenization of parental copies), such as *FBgn0032740* and *Cyp6t1*; 3) for 10 cases excluded by the K-dataset, we examined phylogenetic trees provided by Ensembl and confirmed that new gene are derived as suggested by longer branch length; 4) for 36 genes excluded in the K-dataset, FlyBase and Ensembl do not annotate orthologous genes in the outgroup species for most genes, which are also consistent with the lack of Tblastn hits.

Compared to the K-dataset, the false positives and false negatives are much fewer in the G-dataset (Fig. 1, Supplemental Fig. S1). In the G-dataset, we misidentified only 19 new genes with 14 cases caused by double or triple losses in the outgroups (*e.g. CG2291*). In our pipeline, we first searched against closely-related species and then went for remotely-related species. If at least two independent losses are needed to explain the phylogenetic distribution of orthologs, we will assign a young age to gene *A* by following the maximum parsimony. The underlying assumption is: 1) the possibility of double or triple losses should be low; 2) the genomic alignment between *D. melanogaster* and remotely-related species is less reliable compared to that between *D. melanogaster* and closely-related species. Consistently, we only identified 14 cases with support by at least one additional source (FlyBase, Ensembl). The remaining 5 cases are caused by lack of sensitivity of UCSC genome alignment. Both FlyBase and Ensembl annotate orthologs in the outgroup species, but synteny does not cover the corresponding regions. In opposite, genome alignment built spurious alignment in remotely-related species for 49 new genes identified by Kondo *et al*. However, our Tblastn search could not identify a protein at all. Thus, we referred them as false negatives. All these 68 genes are put into the third category entitled with “Dating problem in this work” (Supplemental Fig. S1).

A fourth category (“Not applicable or difficult for dating”) consists of genes which are most resistant for dating due to their sequence features undesirable for new gene identification. 242 specific new genes claimed by Kondo *et al*. belongs to this category (Supplemental Fig. S1). For 178 out of 242 genes, we found that synteny is in conflict for outgroup species sharing the same phylogenetic relationship relative to *D. melanogaster* (e.g. *D. simulans* and *D. sechellia*). These genes are generally located in repetitive regions (*e.g*. tandem amplification). It is thus likely that the orthologous regions may not be equally well assembled or be subject to speciesspecific gene conversion across these outgroup species. We thus excluded these genes. For example, we masked 19 out of 242 genes including 12 *Ste* genes, 5 Y-linked genes and 2 genes encoded by contigs but not anchored to five major chromosome arms. *Ste* is the X-linked tandem gene families each with redundant copies (Supplemental Fig. S2A). In the UCSC Net track, the most assembles can only reach level 2 of one-way syntenic mapping, rather than the adequate reciprocal syntenic mapping as level 1. The closely related species (*e.g. D. simulans*) also encodes multiple copies, but the corresponding region is not fully assembled and filled with lots of gaps and many of these copies even cannot be assigned to chromosomes (Supplemental Fig. S2B). The size contrast of assemblies between *D. sechellia* and *D. melanogaster* suggests that the region in *D. sechellia* might not be properly assembled due to its repetitive structure (Supplemental Fig. S2A and S2C).

In order to date each member correctly, high-quality outgroup genome must be available first. As for 5 Y-linked genes, Koerich *et al*. (2008) assigned all of them to be old genes (Koerich et al. 2008). The remaining 45 genes consist of three subtypes: 1) 21 fast-evolving small proteins (<100 amino acids); 2) 20 polycistronic genes without previous literature support; 3) 4 tandem duplicates. Different from the aforementioned 178 genes, the syntenic information is consistent across outgroup species suggesting that they are old genes. The small proteins or polycistronic genes are poorly annotated across outgroups. For 4 duplicates, Ensembl phylogenetic tree could not provide diagnostic information to infer the duplication order. Thus, we believed that all these three subtypes are difficult to date as of now. In our private new gene list, 45 candidates also belong to gene families. Similar to the above 4 tandem duplicates, the synteny is consistent across outgroup species showing that these 45 genes are derived copies in their respective families. However, Ensembl phylogenetic tree could not provide additional support. So, we also put these genes into the same fourth category.

The fifth category (“Different ortholog definition”) only consists of 28 private new genes identified by Kondo *et al* (Supplemental Fig. S1). For 24 out of 28 cases, the UCSC syntenic chain only covers a small portion (<30%) of coding regions or mainly corresponds to untranslated regions (UTRs) in outgroup species. Since the dating of Kondo *et al* is protein-centric, they called these genes as new genes. By contrast, our dating pipeline works on DNA-level and identified these genes as old genes. The reason why we took the age of most conserved exons to represent the age of whole genes is that these exons usually represent most important functional regions. Moreover, by performing dating on DNA-level, our dating does not depend on annotation quality of outgroup species. Actually, for all 24 cases, whether the coding region is accurately annotated is unknown due to the lack of protein evidence. For the remaining 4 cases, they all represent translocated genes. In our terminology, we only referred derived duplicate or orphan genes as new genes. By contrast, Kondo *et al*. interpreted incorrectly translocated genes as new genes.

The sixth and final category refers to the aforementioned 12 entries not comparable due to “Gene model change” (Supplemental Fig. S1).

In the main text, we merged “Gene model change”, “boundary cases” and “different ortholog definition” as one dubious (in green color of Fig. 1B) category to simplify.

#### D. melanogaster-specific gene identification

Candidate new genes were initially collected from previous studies (Zhou et al. 2008; Zhang et al. 2010b; Chen et al. 2012). We removed from this list of 233 candidates: 1) any genes whose *D. melanogaster*-specific release 6.05 (http://flybase.org) annotation status is ‘withdrawn’, 2) genes not located on the major chromosome arms 2L, 2R, 3L, 3R, or X, and 3) members of large tandem arrays, including the *Sperm dynein intermediate chain* (Nurminsky et al. 1998; Yeh et al. 2012), *Stellate* (*Ste*), and X: 19,900,000-19,960,000 arrays that are *D. melanogaster*-specific but are impossible to specifically study. We checked syntenic whole-genome alignments of the remaining 84 genes manually using our multi-species alignments at the UCSC Genome Browser. To be conservative, we required that all outgroups including the *D. simulans, D.sechellia, D. yakuba*, and *D. erecta* genome assemblies contained no assembly gaps, transposable elements, or repeats corresponding to the flanking regions of the putative *D. melanogaster*-specific gene. *D. melanogaster*-specific gene origination mechanisms and parental genes were taken from the original studies and confirmed using BLAT and BLASTp. If a gene had multiple significant (e < 10^-10^) full-length BLASTp hits in *D. melanogaster*, the hit that was most similar to the *D. melanogaster-specific* gene was assumed to be the parent. We used available *D. simulans* and *D. yakuba* next-generation sequencing reads to test the presence of putative *D. melanogaster-specific* tandem duplications in these two species (Green et al. 2014; Rogers et al. 2014). We found no breakpoint spanning read pairs supporting *D. melanogaster* tandem duplications in any of 20 *D. simulans* or 20 *D. yakuba* genomes. Thus, these tandem duplications are specifically found in *D. melanogaster* and are not simply missing from the *D. yakuba* and *D. simulans* reference genome assemblies. We checked if any of the duplications in our final set are segregating rather than being fixed within *D. melanogaster* by analyzing 17 whole genome resequencing data from the DPGP2 core Rwanda (RG) genomes (Ni et al. 2008). We required tandem duplications to have at least one read uniquely mapped to each of the three unique breakpoints in order to be called as ‘present’ in a particular line. Ten genes are not found in any of 17 additional *D. melanogaster* genomes we analyzed, suggesting that they are found specifically in the reference stock. Finally, we curated 10 *D. melanogaster*-specific genes. This dataset is actually a subset of G_K new gene data list.

### RNAi strain construction

Since species-specific new genes are under-represented in public RNAi lines, we generated new RNAi lines following Dietzl et al. (2007). Briefly speaking, we designed RNAi reagents using the E-RNAi server (http://www.dkfz.de/signaling/e-rnai3/) and kept constructs with all possible 19-mers uniquely matching the intended target gene and excluded designs with >1 CAN repeat (simple tandem repeats of the trinucleotide with N indicates any base) (Ma et al. 2006). Constructs were cloned into pKC26 following the Vienna *Drosophila* Resource Center’s (VDRC’s) KK library strategy (http://stockcenter.vdrc.at, last accessed 2 February 2016). We introgressed the X chromosome from Bloomington *Drosophila* Stock Center line 34772, which expresses ΦC31 integrase in ovary under control of the *nanos* promoter, into the VDRC 60100 strain. Strain 60100 carries attP sites at 2L:22,019,296 and 2L:9,437,482 (Green et al. 2014). We ensured that our RNAi constructs were inserted only at the 2L:9,437,482 site using PCR following Green et al. (2014). RNAi constructs were injected into the 60100-ΦC31 at 250 ng/μL. Surviving adult flies were crossed to sna^Sco^/CyO balancer flies (BDSC 9325) and individual insertion strains were isolated by backcrossing.

### RNAi screen

We knocked down target gene expression using driver lines constitutively and ubiquitously expressing GAL4 under the control of either the *Actin5C* or *αTubulin84B* promoter. We replaced driver line’s balancer chromosomes with GFP-marked chromosomes to track non-RNAi progeny. Control crosses used flies from the background strains 60100-ΦC31, 25709, or 25710 crossed to driver strains. Five males and five virgin driver females were used in each cross. Crosses were grown at 25°C, 40% - 60% humidity, and a 12h:12h light:dark cycle. F1 progeny were counted at day 19 after crossing, after all pupae had emerged. We screened F1 RNAi flies for visible morphological defects in 1) wings: vein patterning and numbers, wing periphery; 2) notum: general bristle organization and number, structure and smoothness; 3) legs: number of segments. We monitored survival of RNAi F1s by counting GFP and non-GFP L1, L3 larvae and pupae. We tested RNAi F1 sterility by crossing individual RNAi F1 flies to 60100-ΦC31 and monitoring vials for L1 production. Ten replicates for each sex for each line were performed.

### RNAi knockdown specificity and sensitivity

We sought to address two known problems of RNAi technology using RT-qPCR. First, since off-target effects are often discussed in RNAi experiments (Dietzl et al. 2007) we need to test whether target gene expression are specifically knocked down, although our constructs are computationally predicted to be specific. Second, since the RNAi knockdown is often incomplete (Dietzl et al. 2007), we need to estimate how many genes are adequately knockdown in expression. We targeted a random dataset of 14 *D. melanogaster*-specific genes. We collected qPCR primers from FlyPrimerBank (Hu et al. 2013). For those genes not found in FlyPrimerBank, we took Primer-BLAST to design primers by specifically targeting a ~100 bp region of the gene (Supplemental Table S7). We confirmed primer specificity with PCR and Sanger sequencing.

We randomly selected 75 KK RNAi lines (no *tio* site insertion) to analyze their knock down efficiency. We cross these 75 KK RNAi lines with same driver which was used in Dietzl et al., 2007 for GD RNAi line knock down efficiency test. We extracted RNA from sets of 8 adult males (2~4 day old) in triplicate from each RNAi cross using TRIzol (Catalog# 15596-026, Invitrogen, USA), treated ~2 μg RNA with RNase-free DNase I (Catalog# M0303S NEW ENGLAND Biolabs, USA), then used 1 μL treated RNA in cDNA synthesis with SuperScript III Reverse Transciptase (Invitrogen, USA) using oligo(dT)_20_ primers. cDNA was diluted 1:40 in water before using 1 μL as template in 10 μL qPCRs with iTaq™ Universal SYBR Green Supermix (Catalog# 1725121, Bio-Rad, USA) and 400 nM each primer. Reactions were run on a Bio-Rad C1000 Touch thermal cycler with CFX96 detection system (BioRad, CA). Cycling conditions were 95°C for 30 sec, then 45 cycles of 95°C for 5 sec, 60°C for 30 sec, and 72°C for 15 sec. We normalized gene expression levels using the *ΔΔC_T_* method and *RpL32* as the control (Livak and Schmittgen 2001; Dietzl et al. 2007). We tested the specificity and efficiency (90%< qPCR Efficiency<110%) of qPCR primers using an 8-log_2_ dilution series for each primer pair (VanKuren and Long 2018).

### Testing Compensation Effects of New Gene Duplicates

We generated the frameshift mutation line of *vis* using the CRISPR protocol previously developed (VanKuren and Long 2018) but with one single sgRNA for one gene as Kondo et al (2017) did. The sgRNA-*vis* primer below was synthesized (the underlined sequence): 5’-GAAATTAATACGACTCACTATAGGATGTACGGCAGAACATAAGTTTAAGAGCTATGCTGGAA-3’;

We used the following sequence-specific qRT-PCR primers to test the compensatory expression of *achi*, the duplicate of *vis*. Two control genes including *CG12608* and *hth* were examined too. Since *vis*’s expression is largely testis-specific, we extracted RNAs from testis of mated 4-day males and used qRT-PCR with 3 replicates to assess the expression, as developed previously (VanKuren and Long 2018).

248bp:

Achi-RT1F: 5’-AAAGTGACAGGTTTCTCTGTTTG-3’;

Achi-RT1R: 5’-CTGATCCTCCTCCACGATGAC-3’.

237bp:

CG12608-RT1F: 5’-CATAGTGGGCACCTACGAG-3’;

CG12608-RT1R: 5’-TGCGAGAGTATGATCTGCGAC-3’.

92bp:

hth-RT1F:5’-CCTAGTCATGTATCGCCGGTC-3’;

hth-RT1R: 5 ‘-AGCGGATGTTCATAAATC GCA-3’.

Internal control:

113bp:

RpL32-RT1F: 5’-AGCATACAGGCCCAAGATCG-3’;

RpL32-RT1R: 5’-TGTTGTCGATACCCTTGGGC-3’.

## ACKNOWLEDGMENTS

We are grateful for valuable discussion with Edwin Ferguson, Urs Schmidt-Ott, Norbert Perrimon, Emily Mortola, the M. Long lab members and Y.E. Zhang lab members for the technical development in RNAi and scientific issues related to this study. We also appreciate Lisa Meadows from VDRC for supplying haplotyping results for KK lines of the correct insertion site. Y.E. Zhang was supported by the National Key R&D Program of China (2019YFA0802600, 2018YFC1406902) and the National Natural Science Foundation of China (91731302, 31771410, 31970565). M. Long was supported by NSF1026200 and NIH R01GM116113.

## Supplementary Figures

**Figure S1.** Age dating between this work and Kondo *et al*. This figure, following Fig. 1 in the main text, adds specific information on how we classified genes into six major categories or dozens of subcategories. For more details, please refer to Materials and Methods.

**Figure S2.** A representative difficult-to-date locus in the *K*-dataset. A. The syntenic view of *Ste* locus between *D. melanogaster* and *D. simulans* shows fragmented continuity. Due to its multiplicative nature, *Ste* locus is difficult to assemble. In the UCSC Net track, the most assembles can only reach level 2 of one-way syntenic mapping, rather than a better reciprocal syntenic mapping as level 1. B. Some orthologous region in *D. simulans* (lifted from *D. melanogaster*) is not anchored to the chromosome (X) and they are arbitrarily assembled as chrU. C. In *D. sechellia*, two scaffolds are assembled with the major scaffold super_20 spanning 200 kb, in contrast to the assembly of 15 kb for the orthologous region of *D. melanogaster*.

## Supplementary Tables

**Table S1**. The list of genes with the exact ages across the G-dataset and the K-dataset, genes with slightly younger ages in the K-dataset and genes with slightly older ages in the K-dataset, respectively.

**Table S2**. The list of false negatives and false positives in the K-dataset. Since the *Pan-Drosophilid* age group in the K-dataset corresponds to the age group 0, 1 or 2 in the G-dataset (Fig. 1A), we simply replaced the *Pan-Drosophilid* age group as 0/1/2 in the table if applicable.

**Table S3**. 103 knockdown experiments repeated by two independent works (Chen et al. 2010; Zeng et al. 2015). Note, Chen et al (2010) works classified phenotypes as lethal, semi-lethal and viable. Since there are only few genes deemed as semi-lethal, we merged them into lethal gene groups to simplify.

**Table S4**. For 86 new genes with different RNAi drivers, the consistency between different drivers in Chen et al (2010) and Zeng et al (2015) is listed.

**Table S5**. The knockdown efficiency data of KK library and GD library.

**Table S6**. The genotyping results of 153 KK lines, the corrected lines by recombination and knockdown results.

**Table S7**. Primers for 75 KK lines knockdown efficiency tests.

## Notes

### Competing Interest Statement

The authors have declared no competing interest.

### Summary of Updates

Change title

## REFERENCES

1. Ashburner M, Misra S, Roote J, Lewis SE, Blazej R, Davis T, Doyle C, Galle R, George R, Harris N et al. 1999. An exploration of the sequence of a 2.9-Mb region of the genome of Drosophila melanogaster: the Adh region. Genetics 153: 179–219.

2. Carroll SB. 2005. Endless Forms Most Beautiful: The New Science of Evo Devo. W. W. Norton & Company

3. Carvunis AR, Rolland T, Wapinski I, Calderwood MA, Yildirim MA, Simonis N, Charloteaux B, Hidalgo CA, Barbette J, Santhanam B et al. 2012. Protogenes and de novo gene birth. Nature 487: 370–374.

4. Chen SD, Krinsky BH, Long MY. 2013. New genes as drivers of phenotypic evolution. Nat Rev Genet 14: 645–660.

5. Chen SD, Spletter M, Ni XC, White KP, Luo L, Long M. 2012. Frequent recent origination of brain genes shaped the evolution of foraging behavior in Drosophila. Cell Rep 1: 118–132.

6. Chen SD, Zhang YE, Long MY. 2010. New genes in Drosophila quickly become essential. Science 330: 1682–1685.

7. Dietzl G, Chen D, Schnorrer F, Su KC, Barinova Y, Fellner M, Gasser B, Kinsey K, Oppel S, Scheiblauer S et al. 2007. A genome-wide transgenic RNAi library for conditional gene inactivation in Drosophila. Nature 448: 151–156.

8. Ding Y, Zhou Q, Wang W. 2013. Origins of new genes and evolution of their novel functions. Annu Rev Ecol Evol Syst 43: 345–363.

9. El-Brolosy MA, Kontarakis Z, Rossi A, Kuenne C, Gunther S, Fukuda N, Kikhi K, Boezio GLM, Takacs CM, Lai SL et al. 2019. Genetic compensation triggered by mutant mRNA degradation. Nature 568: 193–197.

10. El-Brolosy MA, Stainier DYR. 2017. Genetic compensation: A phenomenon in search of mechanisms. Plos Genet 13: e1006780.

11. Gould SJ. 2002. The structure of evolutionary theory. Kelknap Press of Harvard University Press. Cambridge, Massashusetts and London, England.

12. Green EW, Fedele G, Giorgini F, Kyriacou CP. 2014. A Drosophila RNAi collection is subject to dominant phenotypic effects. Nat Methods 11: 222.

13. Hu Y, Sopko R, Foos M, Kelley C, Flockhart I, Ammeux N, Wang X, Perkins L, Perrimon N, Mohr SE. 2013. FlyPrimerBank: an online database for Drosophila melanogaster gene expression analysis and knockdown evaluation of RNAi reagents. G3 3: 1607–1616.

14. Huang N, Lee I, Marcotte EM, Hurles ME. 2010. Characterising and predicting haploinsufficiency in the human genome. Plos Genet 6: e1001154.

15. Jacob F. 1977. Evolution and tinkering. Science 196: 1161–1166.

16. Jiang X, Assis R. 2017. Natural selection drives rapid functional evolution of young Drosophila duplicate genes. Mol Biol Evol 34:3089–3098.

17. Kasinathan B, Colmenares SU, McConnell H, Young JM, Karpen GH, Malik HS, 2020. Innovation of heterochromatin functions drives rapid evolution of essential ZAD-ZNF genes in *Drosophila*. bioRxiv doi.org/10.1101/2020.07.08.192740.

18. Kersey PJ, Allen JE, Armean I, Boddu S, Bolt BJ, Carvalho-Silva D, Christensen M, Davis P, Falin LJ, Grabmueller C. 2015. Ensembl Genomes 2016: more genomes, more complexity. Nucleic Acids Res 44: D574–D580.

19. Koerich LB, Wang XY, Clark AG, Carvalho AB. 2008. Low conservation of gene content in the Drosophila Y chromosome. Nature 456: 949–951.

20. Kondo S, Vedanayagam J, Mohammed J, Eizadshenass S, Kan LJ, Pang N, Aradhya R, Siepel A, Steinhauer J, Lai EC. 2017. New genes often acquire male-specific functions but rarely become essential in Drosophila. Genes Dev 31: 1841–1846.

21. Krebs JE, Gildstein ES and Kilpatrick ST. 2013. Lewin’s essential genes. Jones & Bartlett Publishers.

22. Lee YCG, Ventura IM, Rice GR, Chen D-Y, Colmenares SU, and Long M. 2019. Rapid Evolution of Gained Essential Developmental Functions of a Young Gene via Interactions with Other Essential Genes. Mol Biol Evol 36: 2212–2226.

23. Liebeskind BJ, McWhite CD, Marcotte EM. 2016. Towards Consensus Gene Ages. Genome Biol Evol 8: 1812–1823.

24. Livak KJ, Schmittgen TD. 2001. Analysis of relative gene expression data using real-time quantitative PCR and the 2(T)(-Delta Delta C) method. Methods 25: 402–408.

25. Long MY, Langley CH. 1993. Natural selection and the origin of jingwei, a chimeric processed functional gene in Drosophila. Science 260: 91–95.

26. Long, MY, Betrán E, Thornton K, and Wang W. 2003. The origin of new genes: glimpses from the young and old. Nature Reviews Genetics 4: 865–875.

27. Long MY, VanKuren NW, Chen SD, Vibranovski MD. 2013. New gene evolution: Little did we know. Annu Rev Genet 47: 307–333.

28. Loppin B, Lepetit D, Dorus S, Couble P, Karr TL. 2005. Origin and neofunctionalization of a Drosophila paternal effect gene essential for zygote viability. Current Biology 15: 87–93.

29. Ma Y, Creanga A, Lum L, Beachy PA. 2006. Prevalence of off-target effects in Drosophila RNA interference screens. Nature 443: 359–363.

30. Ma Z, Zhu P, Shi H, Guo L, Zhang Q, Chen Y, Chen S, Zhang Z, Peng J, Chen J. 2019. PTC-bearing mRNA elicits a genetic compensation response via Upf3a and COMPASS components. Nature 568:259–263.

31. Mayr EJ. 1982. The Growth of Biological Thought-Diversity, Evolution, and Inheritance. New York Rev Books 29: 41–42.

32. Ni JQ, Markstein M, Binari R, Pfeiffer B, Liu LP, Villalta C, Booker M, Perkins L, Perrimon N. 2008. Vector and parameters for targeted transgenic RNA interference in Drosophila melanogaster. Nat Methods 5: 49–51.

33. Nurminsky DI, Nurminskaya MV, De Aguiar D, Hartl DL. 1998. Selective sweep of a newly evolved sperm-specific gene in Drosophila. Nature 396: 572–575.

34. Perrimon N, Engstrom L, Mahowald AP. 1989. Zygotic Lethals with Specific Maternal Effect Phenotypes in Drosophila-Melanogaster .1. Loci on the X-Chromosome. Genetics 121: 333–352.

35. Raices JB, Otto PA, Vibranovski MD. 2019. Haploid selection drives new gene male germline expression. Genome Res 29: 1115–1122.

36. Rhead B, Karolchik D, Kuhn RM, Hinrichs AS, Zweig AS, Fujita PA, Diekhans M, Smith KE, Rosenbloom KR, Raney BJ et al. 2010. The UCSC Genome Browser database: update 2010. Nucleic Acids Res 38: D613–D619.

37. Rogers RL, Cridland JM, Shao L, Hu TT, Andolfatto P, Thornton KR. 2014. Landscape of standing variation for tandem duplications in Drosophila yakuba and Drosophila simulans. Mol Biol Evol 31: 1750–1766.

38. Ross BD, Rosin L, Thomae AW, Hiatt MA, Vermaak D, de la Cruz AFA, Imhof A, Mellone BG, Malik HS. 2013. Stepwise evolution of essential centromere function in a Drosophila neogene. Science 340: 1211–1214.

39. Rossi A, Kontarakis Z, Gerri C, Nolte H, Holper S, Kruger M, Stainier DYR. 2015. Genetic compensation induced by deleterious mutations but not gene knockdowns. Nature 524: 230–235.

40. Ruiz-Orera J, Verdaguer-Grau P, Villanueva-Cañas JL, Messeguer X, Mar Albà M. 2018. Translation of neutrally evolving peptides provides a basis for de novo gene evolution. Nature Ecol Evol 2, 890–896.

41. Schroeder, CM, Tomlin, SA, Valenzuela, JR and Malik, HS, 2020. A rapidly evolving actin mediates fertility and developmental tradeoffs in Drosophila. bioRxiv.

42. Serobyan V, Kontarakis Z, El-Brolosy MA, Welker JM, Tolstenkov O, Saadeldein AM, Retzer N, Gottschalk A, Wehman AM, Stainier DYR, 2020. Transcriptional adaptation in Caenorhabditis elegans eLife 9: e50014.

43. Shao Y, Chen C, Shen H, He BZ, Yu D, Jiang S, Zhao S, Gao Z, Zhu Z, Chen X et al. 2019. GenTree, an integrated resource for analyzing the evolution and function of primate-specific coding genes. Genome Res 29: 682–696.

44. St Pierre SE, Ponting L, Stefancsik R, McQuilton P, FlyBase C. 2014. FlyBase 102--advanced approaches to interrogating FlyBase. Nucleic Acids Res 42: D780–D788.

45. Vakirlis N, Acar O, Hsu B, Castilho Coelho N, Van Oss SB, Wacholder A, Medetgul-Ernar K, Bowman RW, 2nd, Hines CP, Iannotta J et al. 2020. De novo emergence of adaptive membrane proteins from thymine-rich genomic sequences. Nat Commun 11: 781.

46. VanKuren NW, Long MY. 2018. Gene duplicates resolving sexual conflict rapidly evolved essential gametogenesis functions. Nat Ecol Evol 31: 705–712.

47. Ventura IM. 2019. Functional Evolution of Young Retrogenes with Regulatory Roles in Drosophila. The University of Chicago Ph.D. dissertation 10.6082/uchicago.1799.

48. Vissers JHA, Manning SA, Kulkarni A, Harvey KF. 2016. A Drosophila RNAi library modulates Hippo pathway-dependent tissue growth. Nat Commun 7: 10368.

49. Witt E, Benjamin S, Svetec N, Zhao L. 2019. Testis single-cell RNA-seq reveals the dynamics of de novo gene transcription and germline mutational bias in Drosophila. eLife 8: e47138.

50. Xie C, Bekpen C, Kunzel S, Keshavarz M, Krebs-Wheaton R, Skrabar N, Ullrich KK, Tautz D. 2019. A de novo evolved gene in the house mouse regulates female pregnancy cycles. elife 8.

51. Yeh SD, Do T, Chan C, Cordova A, Carranza F, Yamamoto EA, Abbassi M, Gandasetiawan KA, Librado P, Damia E. 2012. Functional evidence that a recently evolved Drosophila sperm-specific gene boosts sperm competition. Proc Natl Acad Sci 109: 2043–2048.

52. Zeng XK, Han LL, Singh SR, Liu HH, Neumuller RA, Yan D, Hu YH, Liu Y, Liu W, Lin XH et al. 2015. Genome-wide RNAi screen identifies networks involved in intestinal stem cell regulation in Drosophila. Cell Rep 10: 12261238.

53. Zhang L, Ren Y, Yang T, Li G, Chen J, Gschwend AR, Yu Y, Hou G, Zi J, Zhou R et al. 2019. Rapid evolution of protein diversity by de novo origination in Oryza. Nat Ecol Evol 3: 679–690.

54. Zhang, YE, Vibranovski, MD, Landback P, Marais GA and Long MY. 2010a. Chromosomal redistribution of male-biased genes in mammalian evolution with two bursts of gene gain on the X chromosome. PLoS Biol 8: e1000494.

55. Zhang YE, Vibranovski MD, Krinsky BH, Long M. 2010b. Age-dependent chromosomal distribution of male-biased genes in Drosophila. Genome Res 20: 1526–1533.

56. Zhou Q, Zhang GJ, Zhang Y, Xu SY, Zhao RP, Zhan ZB, Li X, Ding Y, Yang S, Wang W. 2008. On the origin of new genes in Drosophila. Genome Res 18: 1446–1455.

