## Supplemental Fig. for "Genomic Analyses of New Genes and Their Phenotypic Effects Reveal Rapid Evolution of Essential Functions in Drosophila Development"

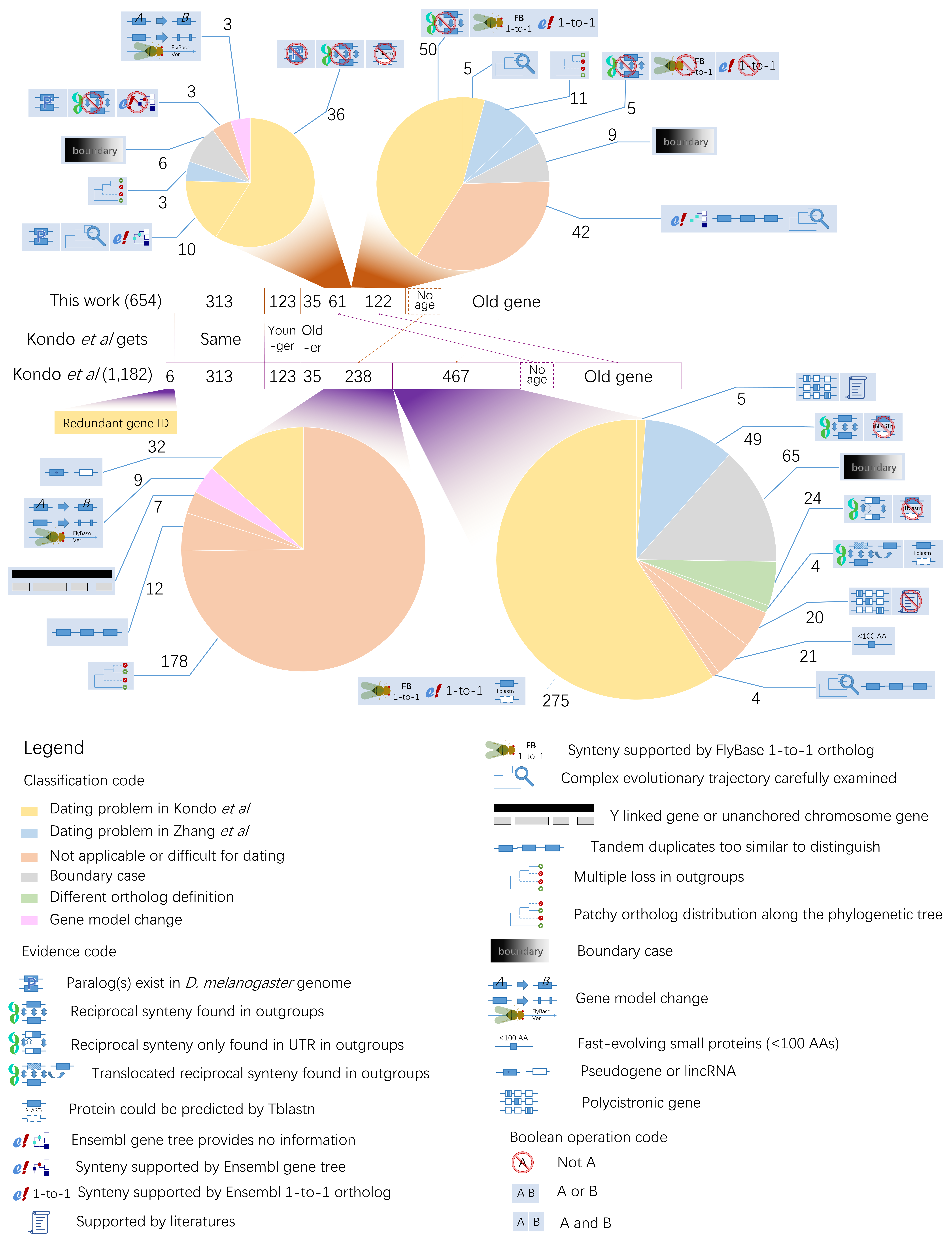

**Figure S1**. Age dating between this work and Kondo et al. This figure, following Fig. 1 in the main text, adds specific information on how we classified genes into six major categories or dozens of subcategories.

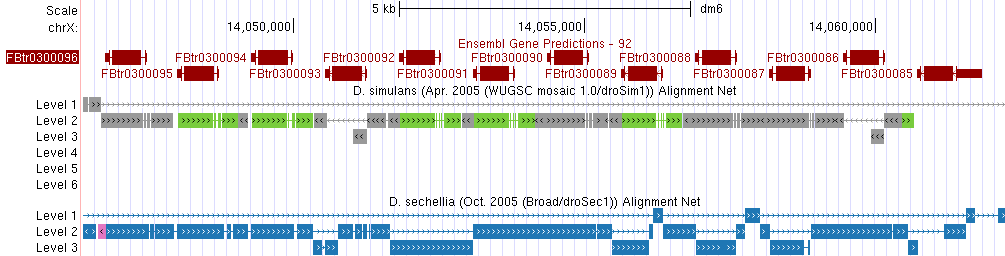
A

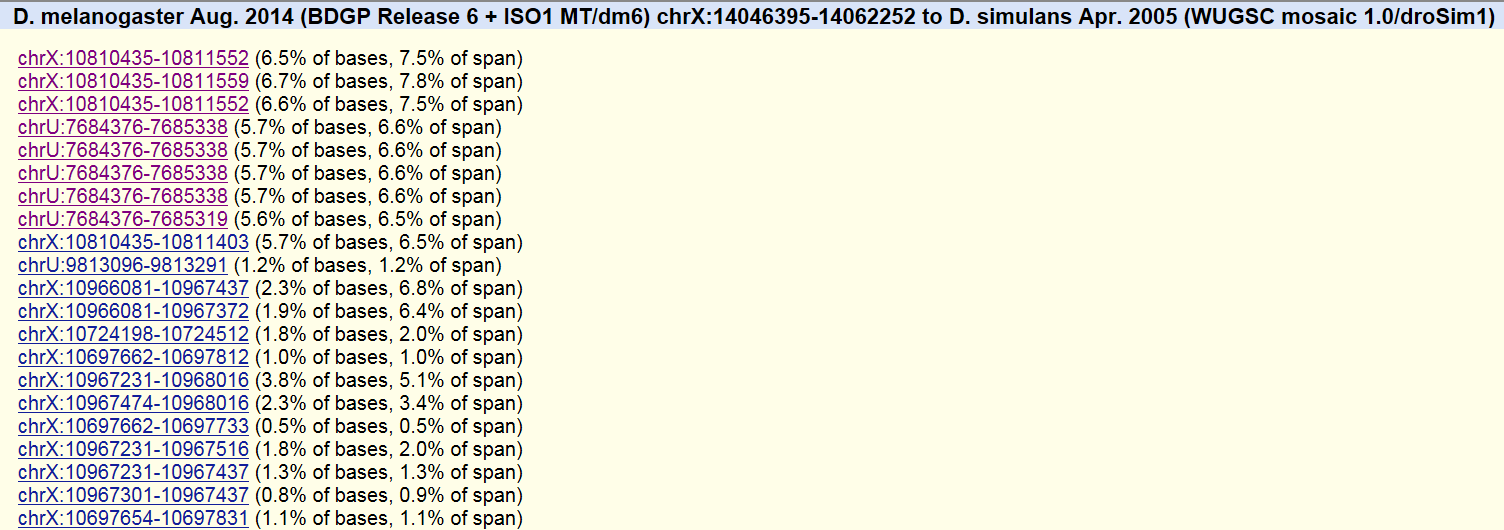
B

C

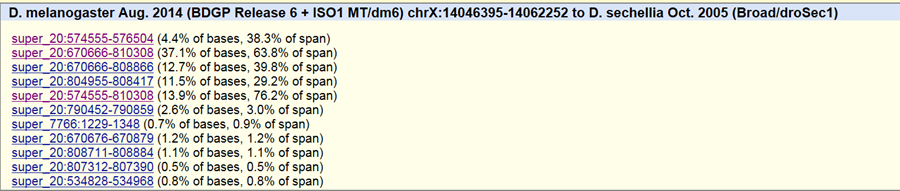

**Figure S2**. **A** representative difficult-to-date locus in the K-dataset. A. The syntenic view of Ste locus between D. melanogaster and D. simulans shows fragmented continuity. Due to its multiplicative nature, Ste locus is difficult to assemble. In the UCSC Net track, the most assembles can only reach level 2 of one-way syntenic mapping, rather than a better reciprocal syntenic mapping as level 1. **B**. Some orthologous region in D. simulans (lifted from D. melanogaster) is not anchored to the chromosome (X) and they are arbitrarily assembled as chrU. **C**. In D. sechellia, two scaffolds are assembled with the major scaffold super_20 spanning 200 kb, in contrast to the assembly of 15 kb for the orthologous region of D. melanogaster.
